# Identification of leaf rust resistance loci in a geographically diverse panel of wheat using genome-wide association analysis

**DOI:** 10.1101/2022.11.10.515255

**Authors:** Shivreet Kaur, Harsimardeep Gill, Matthew Breiland, James Kolmer, Rajeev Gupta, Sunish Sehgal, Upinder Gill

**Author notes:** These authors contributed equally to this work (Shivreet Kaur, Harsimardeep Gill). Corresponding authors: Upinder Gill; Sunish Sehgal.

## Abstract

Leaf rust, caused by *Puccinia triticina (Pt)* is among the most devastating diseases posing a significant threat to global wheat production. The continuously evolving virulent *Pt* races in North America calls for exploring new sources of leaf rust resistance. A diversity panel of 365 bread wheat accessions selected from a worldwide population of landraces and cultivars was evaluated at the seedling stage against four *Pt* races (TDBJQ, TBBGS, MNPSD and, TNBJS). A wide distribution of seedling responses against the four *Pt* races was observed. Majority of the genotypes displayed a susceptible response with only 28 (9.8%), 59 (13.5%), 45 (12.5%), and 29 (8.1%) wheat accessions exhibiting a highly resistant response to TDBJQ, TBBGS, MNPSD and, TNBJS, respectively. Further, we conducted a high-resolution multi-locus genome-wide association study (GWAS) using a set of 302,524 high-quality single nucleotide polymorphisms (SNPs). The GWAS analysis identified 27 marker-trait associations (MTAs) for leaf rust resistance on different wheat chromosomes of which 20 MTAs were found in the vicinity of known *Lr* genes, MTAs, or quantitative traits loci (QTLs) identified in previous studies. The remaining seven significant MTAs identified represent genomic regions that harbor potentially novel genes for leaf rust resistance. Furthermore, the candidate gene analysis for the significant MTAs identified various genes of interest that may be involved in disease resistance. The identified resistant lines and SNPs linked to the QTLs in this study will serve as valuable resources in wheat rust resistance breeding programs.

## Introduction

Global wheat production is continuously constrained by the emergence of new and more virulent races of pathogens causing several economically important diseases. Among these, fungal pathogens are known to cause several important foliar diseases in wheat including three cereal rusts: leaf rust caused by *Puccinia triticina (Pt)*, stem rust caused by *Puccinia graminis* f. sp. *tritici (Pgt)*, and stripe rust caused by *Puccinia striiformis* f. sp. *tritici (Pst)*. Wheat rusts threaten wheat production in the United States (US) by accounting for yield losses in the value of millions of dollars annually. Of the three wheat rusts, leaf rust (LR) is regarded as the most common, extensively distributed, and devastating disease causing 3.25% yield losses annually to global wheat production (Kolmer, 2005; Savary et al., 2019). The recurrent and across-the-board occurrence of leaf rust can lead to epidemic conditions with yield losses ranging from 15% to more than 50% when infections occur during early plant growth stages on susceptible cultivars (Singh et al., 2002; Huerta-Espino et al., 2011). Serious yield losses are incurred in terms of reduced kernels per head and decreased kernel weight (Bolton et al., 2008). In the US alone, ~40-60 races of *Pt* are reported annually (Kolmer et al., 2007) and yield losses valued at $350 million were reported between 2000 and 2004 (Huerta-Espino et al., 2011).

Host resistance is the most efficient and cost-effective strategy to manage leaf rust and wheat breeding programs throughout the world are deploying rust resistance genes in commercial cultivars to fight against this disease (Gill et al., 2019). Around 80 leaf rust resistance (*Lr*) genes have been identified and cataloged in wheat till date (Prasad et al., 2020). Of the identified genes, eleven *Lr* genes have been cloned, viz. *Lr1* (Cloutier et al., 2007), *Lr9* (Wang et al., 2022), *Lr10* (Feuillet et al., 2003), *Lr13* (Hewitt et al., 2021; Yan et al., 2021), *Lr14a* (Kolodziej et al., 2021), *Lr21* (Huang et al., 2003), *Lr22a* (Thind et al., 2017), *Lr34* (Krattinger et al., 2009), *Lr42* (Lin et al., 2022), *Lr58* (Wang et al., 2022), and *Lr67* (Moore et al., 2015). Genetic resistance can be classified into two categories, namely seedling/all-stage resistance (ASR) and adult plant resistance (APR). The seedling resistance is largely qualitative resistance usually controlled by a single major gene, effective at all the developmental stages of the plant life cycle. ASR is associated with a hypersensitive response, a programmed cell death that restricts the pathogen growth and spread. The majority of the studied and characterized leaf rust resistance genes are seedling resistance genes, with *Lr76* (Bansal et al., 2017), *Lr79* (Qureshi et al., 2018) and *Lr80* (Kumar et al., 2021b) being the recent additions to this group. On the other hand, non-race specific, adult-plant resistance (APR) is often partial resistance at the adult plant stage controlled by multiple minor-effect genes with an additive effect and only a sizable proportion of the identified *Lr* genes belong to this category. Among the characterized *Lr* genes, *Lr34* (Dyck, 1987), *Lr46* (Singh et al., 1998), and *Lr67* (Hiebert et al., 2010) are broad-spectrum APR genes, providing partial resistance against all three wheat rusts and powdery mildew (*Blumeria graminis* f. sp. *tritici*). In addition to these genes, *Lr68* (Herrera-Foessel et al., 2012), *Lr74* (Chhetri et al., 2016), *Lr77* (Kolmer et al., 2018b), and *Lr78* (Kolmer et al., 2018a) have also been characterized in hexaploid wheat as APR genes against leaf rust. The race-specific *Lr* genes provide resistance against specific races but the continuously evolving virulent pathogen races render these genes ineffective. Thus, it is important to find novel sources of leaf rust resistance that offer resistance to many different leaf rust races in order to improve the overall durability of resistance in released wheat cultivars.

The recent advancements in sequencing approaches and harmonized community efforts have made genome-wide association studies (GWAS) as an important method for studying the marker-trait associations (Zhu et al., 2008; Tibbs Cortes et al., 2021). In contrast to bi-parental linkage mapping, GWAS extensively utilizes the ancient recombination events that occurred in natural populations for the identification of genomic regions associated with traits of interest. In wheat, GWAS has been used successfully for dissecting the genetic basis of agronomic traits (Sukumaran et al., 2014; Gao et al., 2015; Ward et al., 2019; Sidhu et al., 2020), disease resistance (Gyawali et al., 2018; Phan et al., 2018; Zhu et al., 2020; AlTameemi et al., 2021), quality traits (Kristensen et al., 2018; Chen et al., 2019; Yang et al., 2020) and insect resistance (Joukhadar et al., 2013; Mondal et al., 2016).

In the Great Plains region of the US, leaf rust is the most prevalent rust disease. The hard red spring wheat cultivars grown in the northern plains have resistance genes that include *Lr2a, Lr10, Lr16, Lr21, Lr23*, and *Lr34* (Kolmer, 2019). However, the effectiveness of some of these genes has been reduced due to the continuous emergence of new virulent pathogen races. For example, *Lr21* had been deployed for leaf rust resistance since the mid-2000s and is found in some hard red spring cultivars like Glenn, Faller, and RB07. However, virulent races against this gene have been identified in North Dakota and Minnesota (https://www.ars.usda.gov/midwest-area/stpaul/cereal-disease-lab/docs/lr21-virulence-detected/). Thus, exploring and identifying new resistance sources is highly needed. In the current study, a highly diverse panel of bread wheat accessions from different regions of the world was evaluated for ASR against leaf rust races prevalent in the Northern Great Plains of the US. A high-resolution multi-locus GWAS was also performed to identify genomic regions associated with LR resistance to facilitate the development of resistant wheat cultivars for the future.

## Materials and Methods

### Plant material and *P. triticina* races

We used a diverse panel of 365 hexaploid wheat accessions, including landraces and cultivars from different regions of the world (Supplementary Table S1). The 365 accessions were selected from a larger collection of 890 diverse accessions of hexaploid and tetraploid wheat that was previously resequenced using the sequence capture assay (He et al., 2019). The accessions were obtained from the USDA National Small Grains Collection gene bank and grown for one round of purification and seed increase. The metadata for the 365 accessions can be found in the online repository (http://wheatgenomics.plantpath.ksu.edu/1000EC). Four prevalent *Pt* races (TDBJQ, TBBGS, MNPSD, TNBJS) in the Northern Great Plains of the US were selected for screening the wheat lines at the seedling stage. TBBGS (virulent on genes *Lr1, Lr2a, Lr2c, Lr3, Lr10, Lr21, Lr28*, and *Lr39*) was the most predominant race in Minnesota, North Dakota, and South Dakota in 2020. Furthermore, MNPSD (virulent on genes *Lr1, Lr3, Lr9, Lr24, Lr3ka, Lr17, Lr30, LrB, Lr10, Lr14a*, and *Lr39*), TNBJS (virulent on genes *Lr1, Lr2a, Lr2c, Lr3, Lr9, Lr24, Lr10, Lr14a, Lr21, Lr28*, and *Lr39*) and TDBJQ (virulent on genes *Lr1, Lr2a, Lr2c, Lr3, Lr24, Lr10, Lr14a, Lr21*, and *Lr28*) are the other important races in the region.

### Phenotyping assays for leaf rust seedling screening

The 365 wheat accessions were evaluated in two independent experiments for seedling response to LR. For the leaf rust screening, wheat seedlings at the two-leaf stage were evaluated for their reactions to described races in the biosafety level 2 (BSL 2) facility at Dalrymple Research Greenhouse Complex, North Dakota Agricultural Experiment Station (AES), Fargo. Briefly, five seedlings per each accession along with susceptible checks were used for phenotypic screening for each LR race. Plants were grown in a 50-cell tray containing PRO-MIX LP-15 (www.pthorticulture.com) sterilized soil mix and maintained in a rust-free greenhouse growth room set to 22°C/18°C (day/night) with 16 h/8 h day/night photoperiod. At two-leaf stage, the seedlings were inoculated with fresh urediniospores suspended in SOLTROL-170 mineral oil (Philips Petroleum) at a final concentration of 10^5^ spores mL^-1^ using an inoculator pressurized by an air pump. The inoculated seedlings were placed in a dark dew chamber at 20°C overnight and then transferred back to the growth room. The infection types (IT) were scored about 12 to 14 days after inoculation, using 0-4 scale, where ‘0’ = no visible uredinia, ‘;’ = hypersensitive flecks, ‘1’ = small uredinia with necrosis, ‘2’ = small to medium-sized uredinia with green islands and surrounded by necrosis or chlorosis, ‘3’ = medium-sized uredinia with or without chlorosis, ‘4’ = large uredinia without chlorosis (Stakman et al., 1962). For each IT, ‘+’ or ‘-’ was used to represent variations from the predominant type. A ‘/’ was used for separating the heterogeneous IT scores between leaves with the most prevalent IT listed first. For plants with different ITs within leaves, a range of IT was recorded with the most predominant IT was listed first. The IT scores were converted to a 0-9 linearized scale referred as infection response (IR) (Zhang et al., 2014). Genotypes with linearized IR scores of 0-4 were considered as highly resistant, 5-6 as moderately resistant, and 7-9 as susceptible.

### Phenotypic data analysis

Phenotypic data were analyzed as described previously in (Gill et al., 2021). In brief, a mixed model analysis was used to obtain the best linear unbiased estimates (BLUEs) for phenotypic responses from each of the four isolates using following equation:

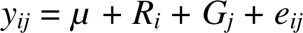

where*y_ij_* is the trait of interest, *μ* is the overall mean, *R_i_* is the effect of the *i^th^* independent replicate/experiment, *G_j_* is the effect of the *j^th^* genotype, and *e_ij_* is the residual error effect associated with the *i^th^* replication, and *j^th^* genotype. The broad-sense heritability (*H^2^*) for IR was estimated for independent nurseries as follows:

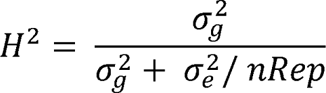

where 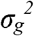 and 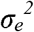 are the genotype and error variance components, respectively. The linear mixed model analysis was performed in META-R (Alvarado et al., 2020) based on the ‘LME4’ R-package (Bates et al., 2015) for the heritability estimation. The Pearson’s correlations among the phenotype responses from four races were estimated based on the BLUEs for each trait using ‘psych’ package in R environment (R Core Team, 2018). The visualization of descriptive statistics was performed using R package ‘ggplot2’ (Wickham, 2016).

### Genotyping, population structure, and linkage disequilibrium

The 365 accessions used in the current study were previously sequenced using the exome sequence capture assay, resulting in the identification of around 7.3 million SNPs (He et al., 2019). A VCF file containing a filtered set of about 3 million SNPs (http://wheatgenomics.plantpath.ksu.edu/1000EC) was used to extract the genotyping information. The extracted genotyping data was subjected to quality control by removing the sites with > 75% missing data, > 5% heterozygotes, and < 5% minimum allele frequency (MAF), leaving 302,524 high-quality SNPs for downstream analysis. The missing genotypes from selected SNPs were imputed by Beagle v4.1 (beagle.27Jan18.7e1.jar; https://faculty.washington.edu/browning/beagle/b4_1.html) (Browning and Browning, 2007) using parameters defined earlier (He et al., 2019) and the imputed set of 302,524 SNPs was used to perform GWAS.

Further, population structure and linkage disequilibrium (LD) analyses were performed using a pruned set of 14,185 SNPs. The LD-based pruning (r^2^ > 0.2) was performed in PLINK v1.90 using ‘indep-pairwise’ function (Purcell et al., 2007). The population stratification was assessed using parallel iterations of a Bayesian model-based clustering algorithms STRUCTURE v2.3.4 (Pritchard et al., 2000; Chhatre and Emerson, 2017) assuming ten fixed populations (*K* =1-10) with ten independent runs for each *K* using a burn-in period of 10,000 iterations followed by 10,000 Monte-Carlo iterations. The optimal value of *K* was identified using STRUCTURE HARVESTER v0.6.9, which is based on an ad-hoc statistic-based approach (Evanno et al., 2005; Earl and vonHoldt, 2012). Further, the principal component analysis (PCA) was performed using 14,185 SNPs with R package ‘SNPRelate’ (Zheng et al., 2012). The first two principal components were plotted as a scatterplot to observe any stratification based on various factors using R package ‘ggplot2’ (Wickham, 2016). The linkage disequilibrium (LD) between SNPs was assessed as the squared correlation coefficient (r^2^) of alleles. The LD decay distance was estimated and visualized for the whole genome and individual sub-genomes using ‘PopLDdecay’ program (Zhang et al., 2019a).

### Marker-trait associations

Genome-wide association analyses were performed using a panel of 365 accessions with 3,02,524 high-quality SNPs to identify marker-trait associations for reaction to four *Pt* isolates (Table S6). Initially, we used two different GWAS algorithms, including the mixed linear model (MLM) (Yu et al., 2006) and a Fixed and random model Circulating Probability Unification (FarmCPU) (Liu et al., 2016). The quantile-quantile (QQ) plots were used to compare the two algorithms that revealed that FarmCPU showed better control of false positives and false negatives. Hence, FarmCPU was used to report GWAS results for all four isolates. In brief, FarmCPU is an improved multiple-locus model that controls false positives by fitting the associated markers detected from the iterations as cofactors to perform marker tests within a fixed-effect model. The FarmCPU was implemented through Genomic Association and Prediction Integrated Tool (GAPIT) version 3.0 in the R environment (Wang and Zhang, 2021), and the first two principal components were included to account for the population structure based upon visual examination of the scree plot and DeltaK statistic from STRUCTURE analysis. The Bonferroni correction-based threshold to declare an association as significant generally proves too stringent as it accounts for all the SNPs in the dataset rather than independent tests. Thus, most studies rely on an exploratory threshold or a corrected Bonferroni threshold based on independent tests (Halder et al., 2019; Pang et al., 2020; Kumar et al., 2021a). In our case, we used an exploratory threshold of −log10(*P*) = 5.00 which is strict compared to the commonly used threshold of −log10(*P*) = 3.00 and suitable for a multi-locus model, which generally does not require multiple corrections (Zhang et al., 2019b). Furthermore, we evaluated the effect of the accumulation of resistant alleles for significant marker-trait associations (MTAs) on the phenotypic performance of the genotypes. The panel of 365 accessions was grouped based on the number of resistant alleles for significant MTAs carried by each accession. These groups were compared using an pairwise t-test to assess the additive effect of the resistant alleles on the disease reaction of respective isolates.

### Candidate gene analysis

The candidate gene analysis was performed for selected stable MTAs to identify the putative candidate genes. As the SNPs were physically mapped to Chinese Spring RefSeq v1.0, we used IWGSC Functional Annotation v1.0 to retrieve high-confidence genes within +/-1Mbp of the significant MTAs. The wheat gene expression browser (http://www.wheat-expression.com/) (Borrill et al., 2016) and a thorough review of the literature was used to exclude unlikely genes from the candidate regions.

## Results

### Phenotypic response of hexaploid wheat accessions to leaf rust

To identify new sources of leaf rust resistance, a panel of 365 hexaploid wheat accessions was phenotypically characterized at the seedling stage with the four *Pt* races. The panel displayed large variations for the disease score ranging from immune response (IT = 0, IR = 0) to highly susceptible response (IT = 4, IR = 9) (Figure 1; Tables 1 and 2). The mean linearized infection response scores of the wheat genotypes were 6.4, 6.3, 5.4, and 6.3 for *Pt* races TDBJQ, TBBGS, MNPSD, and, TNBJS, respectively (Table 1). The distributions of infection responses for all races except MNPSD were skewed towards susceptible scores (IR >7). About 45-50% of the lines displayed susceptible reactions against TDBJQ, TBBGS and, TNBJS, whereas only 12.7% of lines were susceptible to MNPSD with majority of the lines exhibiting a moderately resistance response (Figure 1, Table 2). A total of 28 (9.8%), 59 (13.5%), 45 (12.5%) and 29 (8.1%) wheat lines were highly resistant to TDBJQ, TBBGS, MNPSD and, TNBJS, respectively (Table 2). Majority of the resistant accessions against these *Pt* races were from the Americas followed by Europe, Asia and Africa (Supplementary Table S5). Individually, TDBJQ, TBBGS, MNPSD, and TNBJS had 30.5%, 29%, 28.6%, and 36.7% accessions from the Americas (Supplementary Table S5). Further, there were 71 (19.5%) lines that displayed a resistance response (IR <=6) against all four races (Supplementary Table S6). Out of these 71 resistant accessions, 18.3% were of North American origin (Supplementary Table S6). The proportion of lines showing resistance ranged from 21.4% to 27.9% for the combination of three races and 29.9% to 46.6% for the combination of two races (Table 3). Positive but weak correlations were observed among the seedling plant infection responses to the four *Pt* races and Pearson’s *r* value ranged from 0.16 to 0.39 (Supplementary Figure S1). The broad-sense heritability of infection response for the *Pt* races was high (0.8), showing a large portion of phenotypic variation being explained by the genotypic component (Table 1).

**Figure 1.**
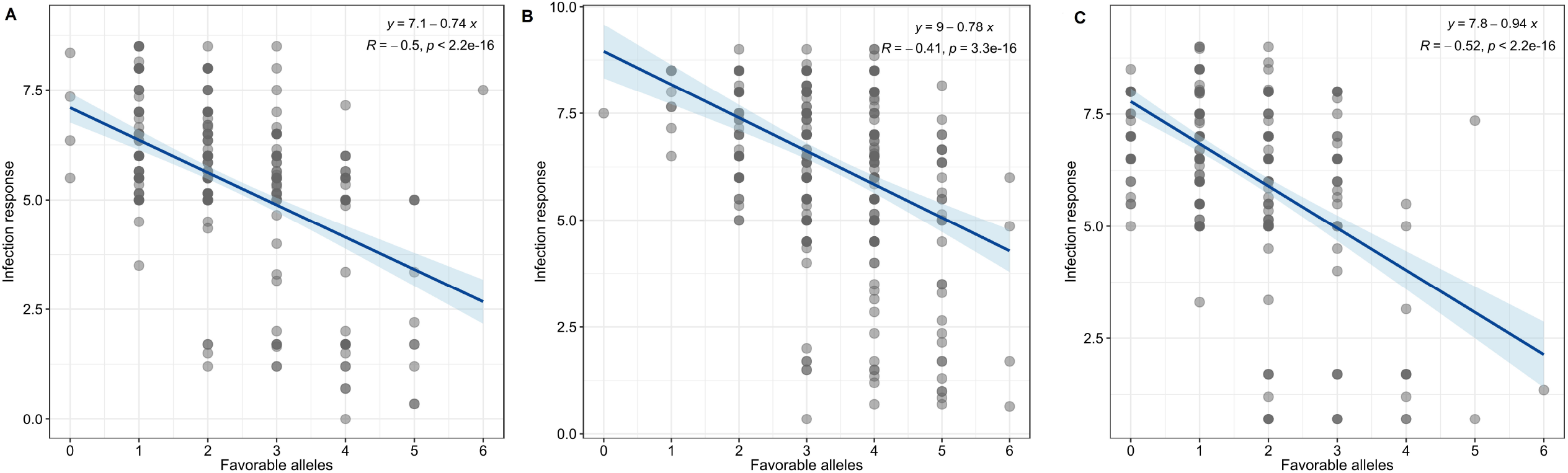
Distribution of infection response (IR) against various races of *Pt* observed during the seedling evaluation of 365 accessions using boxplots and histograms. The X-axis represents the four different *Pt* races and the Y-axis represents the IR in 0-9 scale.

**Table 1.**
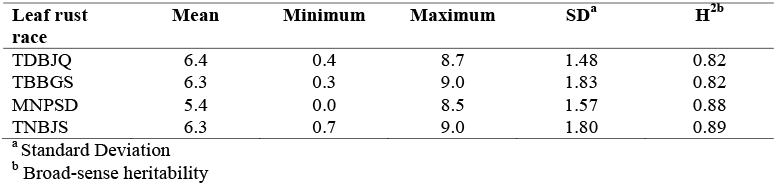
Descriptive statistical analysis of the infection response (IR) calculated from the infection types of the 365 wheat genotypes to *P. triticina* races TDBJQ, TBBGS, MNPSD, and TNBJS.

| Leaf rust race | Mean | Minimum | Maximum | SD<sup>a</sup> | H<sup>2b</sup> |
| --- | --- | --- | --- | --- | --- |
| TDBJQ | 6.4 | 0.4 | 8.7 | 1.48 | 0.82 |
| TBBGS | 6.3 | 0.3 | 9.0 | 1.83 | 0.82 |
| MNPSD | 5.4 | 0.0 | 8.5 | 1.57 | 0.88 |
| TNBJS | 6.3 | 0.7 | 9.0 | 1.80 | 0.89 |

<sup>a</sup> Standard Deviation
<sup>b</sup> Broad-sense heritability

**Table 2.**
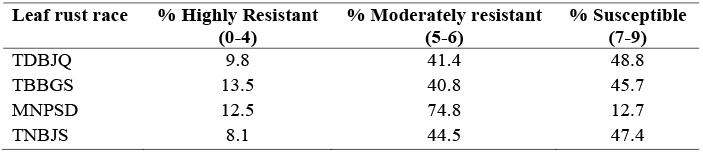
Percent distribution of the diversity panel accessions based on their linearized seedling infection responses (IRs) against *P. triticina* races TDBJQ, TBBGS, MNPSD, and TNBJS. Range of IR score for each category is given in the parenthesis.

| Leaf rust race | % Highly Resistant<br>(0-4) | % Moderately resistant<br>(5-6) | % Susceptible<br>(7-9) |
| --- | --- | --- | --- |
| TDBJQ | 9.8 | 41.4 | 48.8 |
| TBBGS | 13.5 | 40.8 | 45.7 |
| MNPSD | 12.5 | 74.8 | 12.7 |
| TNBJS | 8.1 | 44.5 | 47.4 |

**Table 3.** Number and percentage of lines resistant to different combinations of the four *P. triticina* races.

| Leaf rust race combination | Number of lines<br>(R/MR) | Percentage of lines |
| --- | --- | --- |
| TDBJQ+TBBGS+MNPSD+TNBJS | 71 | 19.5 |
| TDBJQ+TBBGS+MNPSD | 94 | 25.8 |
| TDBJQ+MNPSD+TNBJS | 101 | 27.7 |
| TDBJQ+TBBGS+TNBJS | 78 | 21.4 |
| MNPSD+TBBGS+TNBJS | 102 | 27.9 |
| TDBJQ+TBBGS | 109 | 29.9 |
| TDBJQ+MNPSD | 160 | 43.8 |
| TDBJQ+TNBJS | 112 | 30.7 |
| MNPSD+TBBGS | 170 | 46.6 |
| TBBGS+TNBJS | 111 | 30.4 |
| MNPSD+TNBJS | 167 | 45.8 |

### Population structure and linkage disequilibrium analysis

The STRUCTURE analysis used to infer the population structure revealed two major subpopulations (P1 and P2 hereafter) in the panel of 365 accessions based on the peak of DeltaK statistic (Figure 2a). The subpopulation P1 comprised 82 accessions while P2 was comparatively larger comprising the remaining 283 accessions. Further, we assessed a relationship between the two subpopulations and various characteristics including geographic origin, type, and growth habit of the studied accessions. The subpopulation P1 mostly represented the spring wheat accessions as 78 of the 82 accessions had spring growth habit. In contrast, the subpopulation P2 comprised accessions with both spring (216), winter (53), and facultative (14) growth habits. Additionally, P1 mainly represented landraces (51) with comparatively few cultivars (15); whereas, P2 includes cultivated accessions with 144 cultivars and 69 landraces. Based on geographical origin, P1 includes accessions from Asia (45) and Africa (23) with a few accessions from Europe (3) and the Americas (7). In contrast, P2 includes majority of accessions from Europe (89) and the Americas (95) and a good number of accessions from Asia (40) and Africa (37).

**Figure 2.**
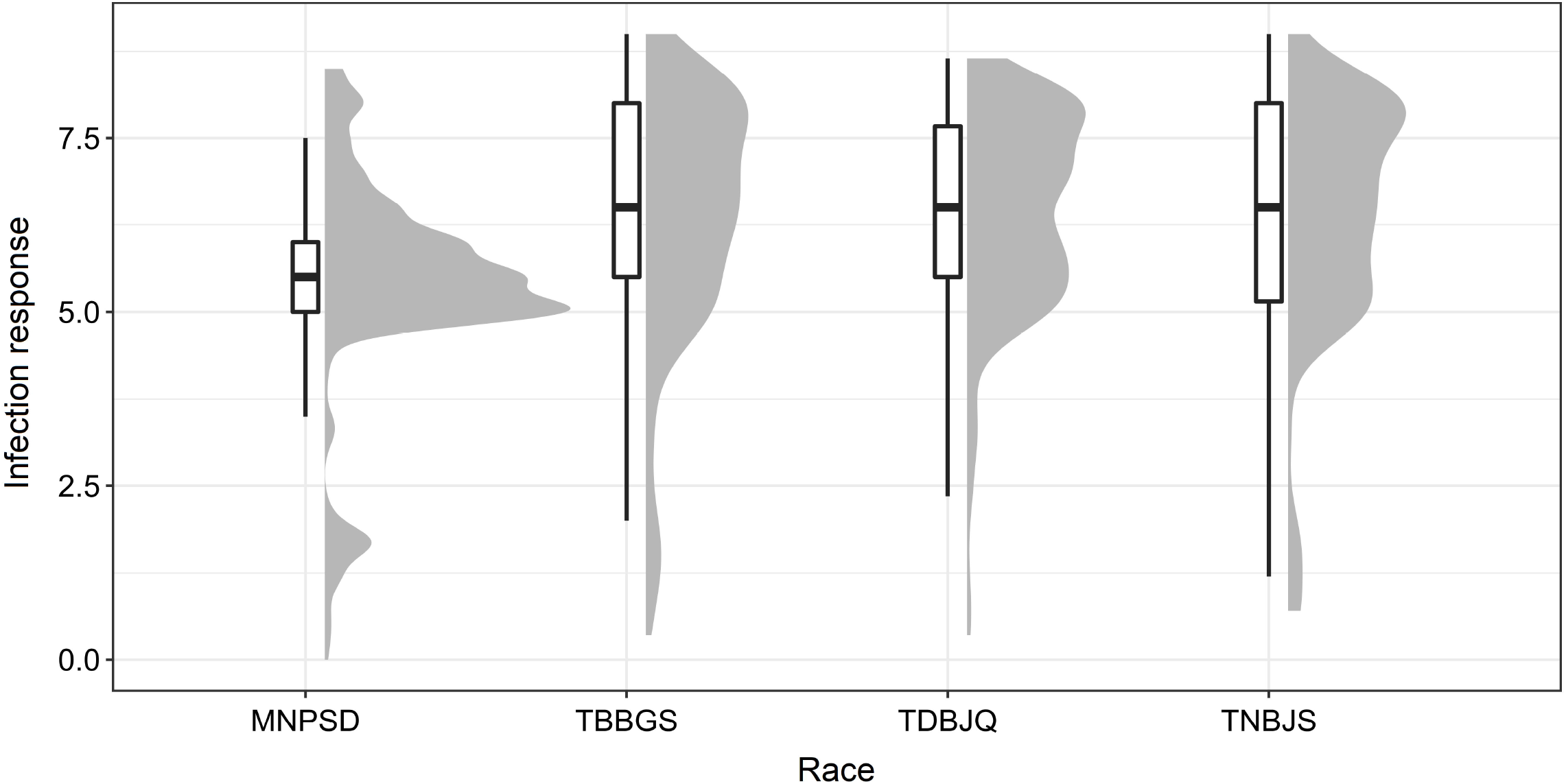
Population structure analysis in panel of 365 wheat accessions based on the 3,02,524 SNPs. (**A**) Evanno plot of Delta-K statistic from the STRUCTURE analysis. (**B**) Scree plot for first 10 components obtained from principal component analysis (PCA). (**C**) Scatterplot based on the first two components (PC1 and PC2) from PCA.

The principal component analysis also revealed two subpopulations within the studied germplasm, with the first two principal components explaining a genetic variation of 4.7% and 3.3%, respectively (Figures 2b and 2c). The first two principal components were plotted to visually differentiate the accessions by origin and growth habit. The PCA results showed a differentiation among the accessions belonging to landrace or cultivar categories (Figure 2c). However, we did not observe a clear differentiation based on growth habit and geographic origin (Supplementary Figure S2). The linkage disequilibrium (LD) decay was estimated based on LD value (*r^2^*) for the whole genome as well as individual sub-genomes. For the whole genome, the LD decay was found to be about 1.5 Mb (Supplementary Figure S3). The LD decay for the three sub-genomes A, B, and D revealed different patterns, with A and B showing smaller LD decay distances as compared to the D sub-genome (Supplementary Figure S3).

### Genome-Wide Association Analyses for leaf rust resistance

The phenotypic data of *Pt* screening was subjected to GWAS to identify genomic regions and SNP markers associated with leaf rust resistance. The GWAS was performed using BLUEs calculated from disease scores data for individual *Pt* races. We identified a total of 27 significant MTAs on twelve chromosomes: 1A, 1B, 2A, 2B, 2D, 3B, 4A, 4B, 4D, 5A, 5B, and 6B for response against the four *Pt* races. Individually, we detected nine, nine, one, and eight significant MTAs for responses against races MNPSD, TBBGS, TDBJQ, and TNBJS, respectively (Figure 3, Table 4). The nine MTAs for race MNPSD were identified on eight different chromosomes including 1A, 2B, 2D, 3B, 4A, 4B, 5A, and 5B (Figure 3, Table 4). The most significant MTA for MNPSD (*scaffold9496_550027;* −log_10_*P* = 11.5) was observed on chromosome 3B physically mapped to 456 Mb and had a SNP effect of 0.91 (Table 4). For response against TBBGS, a total of nine MTAs were identified on seven different chromosomes: 1B, 2A, 3B, 4B, 5A, 5B, and 6B (Figure 3, Table 4). Of note, the MTAs *scaffold145719_3415472, scaffold20863_2950181*, and *scaffold81142-6_3121151* identified on chromosome 1B, 2A, and 5B showed a significant effect value of −0.81, 0.66, and −0.80, respectively. Further, eight significant MTAs were detected for TNBJS mapped on six different chromosomes including 1A, 1B, 2B, 2D, 3B, and 4A (Figure 3, Table 4). Among these eight MTAs, the most significant MTA (*scaffold63719_1362898*) was identified on chromosome 4A at 625 Mb and had a SNP effect value of 0.81. In contrast to the other races, we identified only one MTA (*scaffold38811_1402219*) for isolate TDBJQ mapped at 503 Mb on chromosome 4D (Figure 3, Table 4).

**Figure 3.**
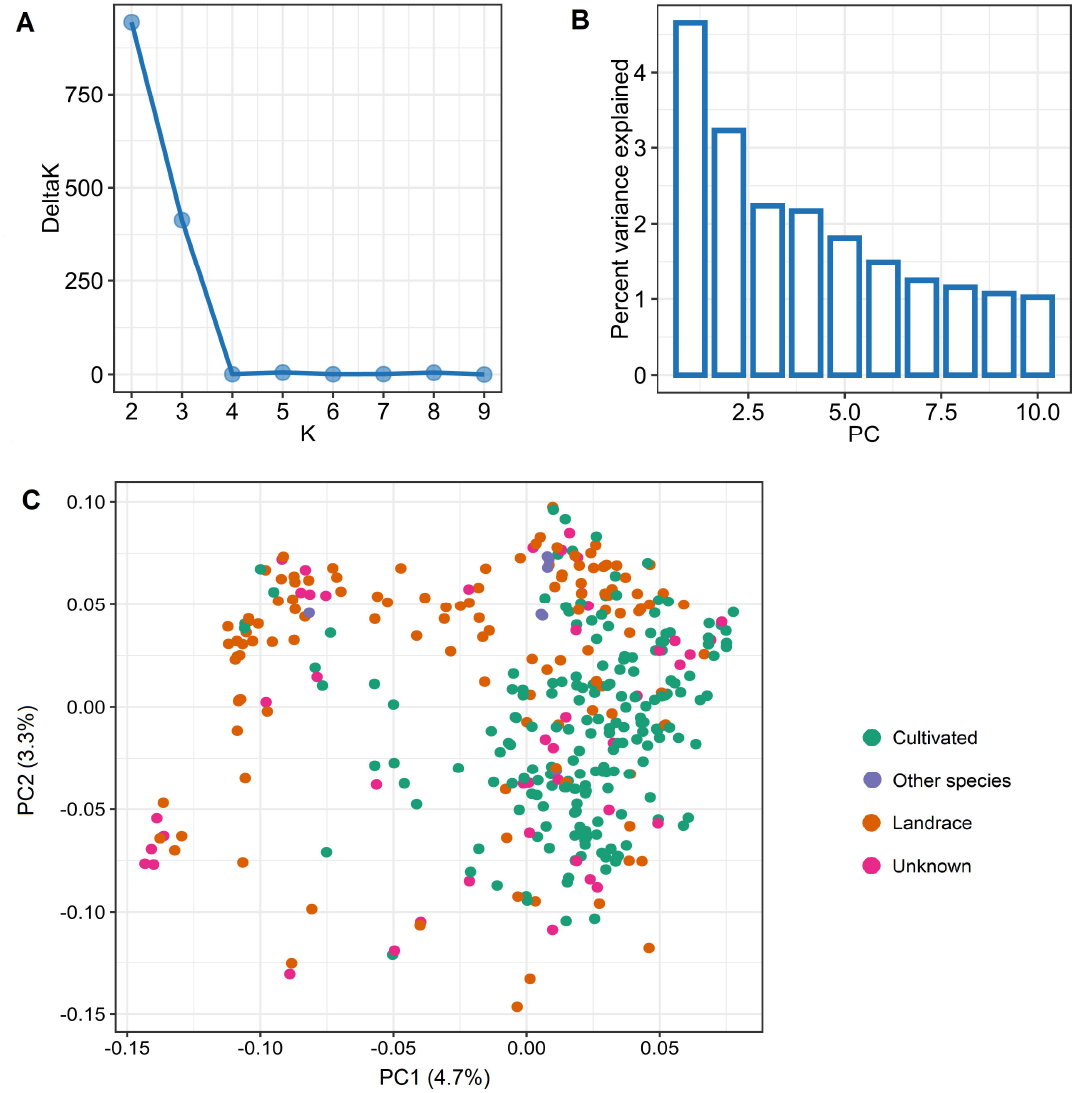
Manhattan plots from genome-wide association studies showing the distinct peaks for identified MTAs in response to (**A**) MNPSD, (**B**) TBBGS, (**C**) TDBJQ, and (**D**) TNBJS races of *P. triticina*. The red horizontal line represents the threshold used to report MTAs for each race.

**Table 4.** Summary of the marker-trait associations (MTAs) identified for resistance to *P. triticina* races MNPSD, TBBGS, TsDBJQ, and TNBJS.

| Race | SNP | Chrom | Position<sup>a</sup> | Allele<sup>b</sup> | P value | Effect<sup>c</sup> | -Log<sub>10</sub>(P) |
| --- | --- | --- | --- | --- | --- | --- | --- |
| MNPSD | scaffold43155_283689 | 1A | 32,381,168 | T/C | 3.9E-07 | 0.36 | 6.4 |
|  | scaffold57495_4340875 | 2B | 448,878,050 | G/T | 5.9E-08 | -0.74 | 7.2 |
|  | scaffold109282_20590799 | 2D | 62,620,318 | C/T | 4.4E-09 | -0.41 | 8.4 |
|  | scaffold9496_550027 | 3B | 456,405,981 | T/C | 3.2E-12 | 0.91 | 11.5 |
|  | scaffold35818-1_1947539 | 4A | 601,155,551 | T/C | 4.8E-06 | 0.28 | 5.3 |
|  | scaffold6836_258869 | 4B | 1,789,216 | G/A | 8.9E-07 | 0.32 | 6.0 |
|  | scaffold31373_401257 | 5A | 30,099,694 | G/C | 1.3E-06 | 0.41 | 5.9 |
|  | scaffold33098_3932746 | 5B | 347,928,790 | T/C | 1.1E-08 | 0.43 | 7.9 |
|  | scaffold81325_356781 | 5B | 586,845,078 | C/T | 9.0E-06 | 0.45 | 5.0 |
| TBBGS | scaffold35219_1114450 | 1B | 17,885,002 | G/T | 3.4E-06 | 0.47 | 5.5 |
|  | scaffold145719_3415472 | 1B | 69,906,201 | C/T | 6.8E-08 | -0.81 | 7.2 |
|  | scaffold33664_2059905 | 2A | 4,217,825 | A/T | 7.6E-07 | 0.40 | 6.1 |
|  | scaffold57658-1_2072073 | 2A | 547,134,074 | T/C | 7.0E-08 | -0.47 | 7.1 |
|  | scaffold22480_722430 | 3B | 544,111,954 | G/A | 4.2E-08 | 0.46 | 7.4 |
|  | scaffold73828-3_704067 | 4B | 538,266,707 | A/G | 3.5E-06 | 0.33 | 5.5 |
|  | scaffold20863_2950181 | 5A | 514,152,049 | T/A | 3.4E-06 | 0.66 | 5.5 |
|  | scaffold81142-6_3121151 | 5B | 115,397,685 | G/T | 2.2E-08 | -0.80 | 7.7 |
|  | scaffold84762_1725496 | 6B | 21,911,761 | A/G | 6.9E-07 | -0.59 | 6.1 |
| TDBJQ | scaffold38811_1402219 | 4D | 503,867,312 | T/C | 1.0E-05 | 0.70 | 5.0 |
| TNBJS | scaffold123808_1588140 | 1A | 21,616,147 | A/G | 1.0E-05 | -0.48 | 5.0 |
|  | scaffold33401_3398330 | 1B | 119,884,884 | G/A | 5.0E-06 | 0.63 | 5.3 |
|  | scaffold56230_857271 | 2B | 671,745,210 | C/T | 1.4E-07 | 0.28 | 6.8 |
|  | scaffold98508-7_1879233 | 2B | 768,104,692 | T/C | 2.0E-08 | 0.34 | 7.7 |
|  | scaffold67556_44513 | 2D | 37,367,713 | T/C | 4.7E-06 | 0.54 | 5.3 |
|  | scaffold151621_1347043 | 3B | 829,196,566 | C/T | 1.1E-08 | -0.79 | 7.9 |
|  | scaffold63719_1362898 | 4A | 625,284,220 | C/T | 8.7E-11 | 0.81 | 10.1 |
|  | scaffold163140-5_613860 | 4A | 725,750,499 | A/G | 6.1E-06 | -0.29 | 5.2 |

<sup>a</sup> Position is based on IWGSC RefSeq v1.0 (IWGSD, 2018)
<sup>b</sup> Allele nomenclature: Major allele/Minor allele
<sup>c</sup> SNP effect

Furthermore, we evaluated the additive effect of resistant alleles of significant MTAs on mean infection response from individual races. As we identified only one MTA for TDBJQ, data from only three races were used for this analysis. Overall, we observed a significant negative association between the number of resistant alleles and infection response for all three races suggesting that the accumulation of resistant alleles in genotypes reduces infection response (Figure 4). The accessions with two or more resistant alleles exhibited a lower infection response against all individual races.

**Figure 4.**
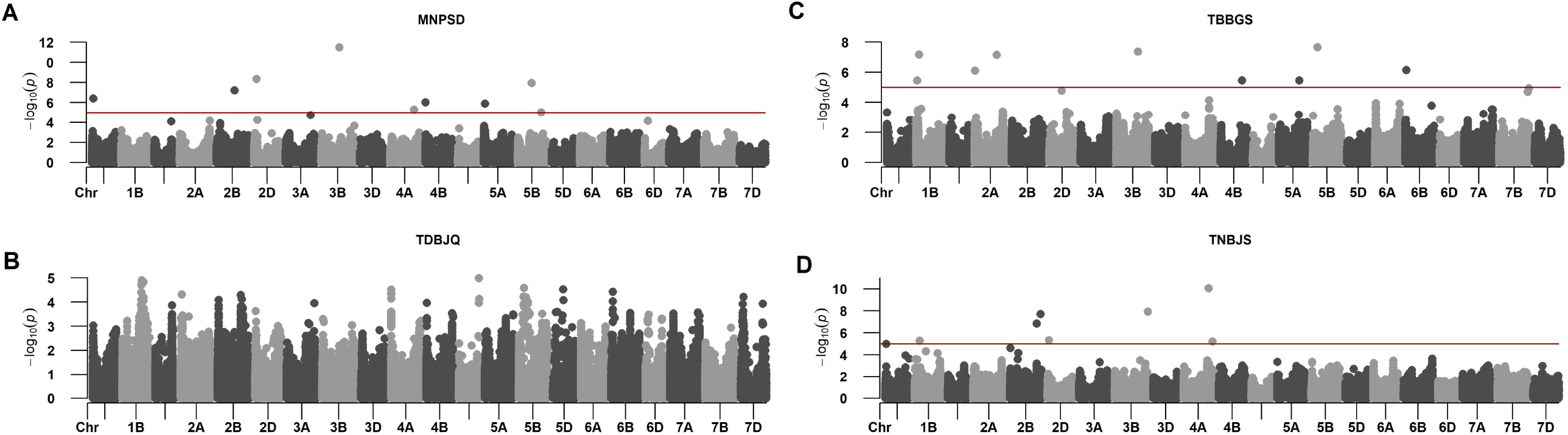
Linear regression plots of seedling response toward *P. triticina* races (**A**) MNPSD, (**B**) TBBGS and, (**C**) TNBJS to the number of favorable alleles of identified MTAs for respective races.

### Candidate gene analysis for significant marker-trait associations

The significant MTAs were further analyzed to identify candidate genes underneath or in close vicinity of the MTAs in the wheat genome. Among the 27 MTAs, 20 were mapped in the proximity of previously reported MTAs, QTLs or genes for LR resistance, validating the importance of these regions. Thus, we selected these regions for identifying putative genes of interest that may play a role in LR resistance. Further, we investigated the local LD decay in these 20 regions by constructing haplotype blocks. Out of the 20 regions, LD decay rate was found to be less than 1Mbp in 17 regions (data not shown). Henceforth, a 1Mbp window around the associated SNP was used to retrieve high-confidence genes using CS RefSeq v1.1. In total, we identified 252 high confidence (HC) genes in +/-1 Mb of the significant MTAs (Supplementary Table S8). Further, a thorough comparison with literature identified genes of interest that encode known plant disease resistance proteins such as intracellular nucleotide-binding and leucine-rich repeat (NLR) receptors, proteins with kinase domains, ATP-binding cassette (ABC) transporters, F-box-like domain-containing proteins, defensins, receptor kinase proteins, and others (Supplementary Table S9).

## Discussion

Wheat germplasm collections serve as major resources for enriching the wheat genetic diversity against various biotic and abiotic stresses including pathogens such as rusts. Utilizing the untapped value of these genetic repositories can help identify potential novel sources of genes and QTLs and thereby advance the course of broadening the resistance diversity against leaf rust. Association studies have been successfully deployed in both bread wheat and durum wheat for detecting genomic regions associated with leaf rust resistance at seedling and adult plant stages (Aoun et al., 2016, 2021; Gao et al., 2016; Li et al., 2016; Sapkota et al., 2019; Leonova et al., 2020). Foliar diseases of wheat like leaf rust are among the most important and destructive diseases in the Northern Great Plains of the US. Identifying novel sources of resistance against leaf rust is a continuing challenge due to the constant evolution of *Pt* populations causing the resistance breakdown of the existing *Lr* genes. In 2020, 15 virulence phenotypes were identified from 140 isolates in North Dakota, Minnesota, and South Dakota with TBBGS (42.1%), MNPSD (17.9%), and TNBJS (5.7%) being the most predominant phenotypes (https://www.ars.usda.gov/midwest-area/stpaul/cereal-disease-lab/docs/cereal-rusts/race-surveys/). Therefore, we evaluated a global wheat diversity panel of 365 hexaploid wheat accessions for identifying genetic loci harboring novel resistance genes for leaf rust against four important and prevalent races namely, TDBJQ, TBBGS, MNPSD, and TNBJS, at the seedling stage.

Our phenotypic evaluations identified a substantial number of resistant accessions from both North and South America (Supplementary Table S5) against the *Pt* races. Further, we identified 59 spring, 10 winter, and 2 facultative growth habit accessions out of the 71 resistant lines carrying race-specific resistance to all four *Pt* races that are prevalent in the north-central region of the United States (Supplementary Table S6). The majority of the accessions are cultivated lines from the Americas (38%) followed by Europe (25.3%), Asia (22.5%), and Africa (9.9%) (Supplementary Table S6). Leaf rust resistance in a sizeable proportion (18.3%) of North American lines could be due to their selection against *Pt* races predominant in North America. Similar results were reported by (Gao et al., 2016), where high resistance was observed in a sub-population consisting mostly North American lines when screened against multiple *Pt* races common in that region. The resistance response to *Pt* races, TNBJS and TBBGS showed a slightly higher correlation (0.39) (Supplementary Figure S1) as compared to the other races which could be due to the similarities in the virulence profile as both races are virulent on *Lr1, Lr2a, Lr2c, Lr3, Lr10, Lr21, Lr28*, and *Lr39*. Out of the 71 resistant lines, 10 wheat accessions displayed a highly resistant response (IR<= 4) against a combination of three different races, and only 3 lines were highly resistant (IR <= 4) against all four races (Supplementary Table S6). These lines may contain one or more existing or novel genes and can serve as a potential source for transferring this resistance into wheat cultivars by the breeding programs.

GWAS with FarmCPU algorithm identified 27 MTAs for leaf rust resistance on the eleven wheat chromosomes (Table 4). The BLUEs were not normally distributed, so we tried BoxCox transformation to normalize the data. However, the transformation did not improve the distribution of BLUEs (Supplementary Figure S4), hence we used non-transformed data for performing GWAS. Out of 27 MTAs, 20 were identified in the vicinity of previously reported genes, MTAs, or QTLs associated with leaf rust resistance. An important MTA (*scaffold33664_2059905*) was identified for LR resistance against TBBGS which mapped around 4 Mb on the short arm of chromosome 2A. Interestingly, this region (0.5 – 7.8 Mb) has been reported to harbor three known *Lr* genes including *Lr17* (Dyck and Kerber, 1977), *Lr37* (Bariana and McIntosh, 1993; Blaszczyk et al., 2004), and *Lr65* (Wang et al., 2010; Mohler et al., 2012; Zhang et al. 2021a), and a recently identified QTL for LR resistance (Fatima et al., 2020) (Supplementary Table S7). A long chromosomal fragment of 25-38 cM containing *Lr37, Yr17*, and *Sr38* was introduced from a wild wheat relative *Aegilops ventricosa* and located on a 2NS/2AS translocation in a winter wheat cultivar ‘VPM1’ (Bariana and McIntosh, 1993). *Lr37* is an adult plant resistance gene but expresses a seedling resistance response of 2+ at temperatures below 20 °C (Park and McIntosh, 1994; Kolmer, 1996). Recently this segment was also found to provide resistance against wheat blast. In addition to *Lr37*, this MTA (*scaffold33664_2059905*)colocalized with *WMS636* (4.9 Mb), a distal flanking marker associated with genes *Lr17a* (Bremenkamp□Barrett et al., 2008) and the two makers, *AltID-11*(0.55Mb) and *Alt-92* (0.61Mb), flanking *Lr65* (Zhang et al. 2021b). Since TBBGS is avirulent against *Lr65* and the identified MTA colocalized with markers tightly linked to *Lr65*, it is highly likely that the locus identified in this study is identical to the *Lr65* locus. Zhang et al. (2021b) identified *TraesCS2A02G001500* encoding NB-ARC and LRR domains as the most prominent candidate gene for *Lr65*. On chromosome 4B, we identified an MTA (*scaffold73828-3_704067*) associated with response to TBBGS in the vicinity of two known *Lr* genes, *Lr12* (Dyck et al., 1966) and *Lr31* (Singh and McIntosh, 1984)*. Lr12* is a race specific adult plant resistance gene and is completely linked or identical to *Lr31* which is a seedling resistance gene that requires another complementary gene *Lr27/Sr2/Yr30* to function (Singh and McIntosh, 1984; Singh et al., 1999; Mago et al., 2011; Singh and Bowden, 2011). However, we did not identify any association in the 3BS region harboring *Lr27* in this study.

Three MTAs (*scaffold35219_1114450, scaffold145719_3415472* and *scaffold33401_3398330*)were identified on chromosome 1B against TBBGS and TNBJS. Two known *Lr* genes, *Lr33* (Dyck, 1987) and *Lr75* (Singla et al., 2017) were previously mapped on chromosome 1B. *Lr75* is an adult plant resistance gene and was first identified in the Swiss cultivar ‘Forno’ (Schnurbusch et al., 2004; Singla et al., 2017). The physical position of *scaffold145719_3415472* approximately co-localized with the physical location of *GWM604*, a distal flanking marker linked to *Lr75* on CS RefSeq v1.0 (Singla et al., 2017). Given that the *scaffold145719_3415472* was identified for seedling resistance to LR, it is unlikely that this MTA represents *Lr75*.Another MTA on chromosome 1B (*scaffold33401_3398330*) was mapped 9.2 Mb apart from a diagnostic marker (*BOBWHITE_C39153_131*) linked to *Lr33* (Che et al., 2019). Interestingly, *scaffold35219_1114450, scaffold145719_3415472* and *scaffold33401_3398330* were mapped within 1 Mb of several previously reported MTAs (*IAAV8117, BS00083533_51*, and *BS00084722_51*) for leaf and stripe rust resistance (Zhang et al., 2021a) validating their role in response to *Pt* (Supplementary Table S7).

We identified two MTAs (*scaffold56230_857271* and *scaffold98508-7_1879233*) for race TNBJS on chromosome 2BL close to *WMS382*, a marker associated with *Lr50* (Brown-Guedira et al., 2003). *Lr50* was transferred from wild wheat, *T. timopheevi armeniacum* and was previously mapped on chromosome 2BL (Brown-Guedira et al., 2003). Furthermore, *scaffold56230_857271*, and *scaffold98508-7_1879233* on 2BL mapped in close vicinity of previously reported MTAs (*AX-95006189* and *AX-94481202*) for LR resistance (Kumar et al., 2020). Another MTA, *scaffold151621_1347043* was mapped in the vicinity of a recently reported MTA (*AX-94671785*) for LR resistance (Vikas et al., 2022) (Supplementary Table S7).

In response to MNPSD, we detected an MTA (*scaffold109282_20590799*) which mapped on chromosome 2D near *aWPT-0330*, a marker linked to a seedling resistance gene, *Lr2* (Tsilo et al., 2014; Dyck and Samborski, 1974). Similarly, *scaffold81325_356781* on chromosome 5B mapped ~ 8 Mbp apart from marker *BOBWHITE_REP_C50349_139* (Carpenter et al., 2017), which is linked to *Lr18* (Dyck and Samborski, 1968). Further studies are needed to determine the relationship between the identified MTAs and the postulated *Lr* genes in close proximity to MTAs. Joukhadar et al. (2020) conducted a GWAS study using 2,300 hexaploid wheat genotypes including worldwide landraces, cultivars and synthetic backcross derivatives for adult plant resistance to leaf rust, stripe and stem rust across multiple Australian environments. Of the 365 wheat accessions used in our study, 257 genotypes overlapped with the adult plant screening panel (Joukhadar et al., 2020); interestingly no common resistant lines and significant marker-trait associations for leaf rust were found among the two studies.

In addition, our analyses also identified two novel MTAs (*scaffold9496_550027* and *scaffold57658-1_2072073*) that were located in genomic regions where no previously reported *Lr* genes or MTAs have been reported in wheat (Supplementary Table S7). Further, we also identified five putatively novel MTAs (*scaffold33098_3932746, scaffold57495_4340875, scaffold163140-5_613860, scaffold151621_1347043*, and *scaffold56230_857271*) mapped at >10 Mb apart from the previously reported MTAs in various GWAS studies (Supplementary Table S7). A total of three, one, and three significant novel MTAs were identified for resistance against MNPSD, TBBGS, and TNBJS, respectively. One of these seven MTAs, *scaffold57495_4340875*, detected on chromosome 2B was found near *Lr35*, (Kerber and Dyck, 1990) an APR gene expressing at two-leaf stage. *BCD260* (Seyfarth et al., 1999), a marker linked to the *Lr35*, was about 30 Mb apart *scaffold57495_4340875*. Given *Lr35* confers APR and the associated marker is far away from the identified MTA, it is unlikely the same locus. Another MTA, *scaffold163140-5_613860* against race TNBJS was mapped on the region harboring known gene *Lr28* (McIntosh et al. 1982) on chromosome 4A. However, TNBJS is virulent on *Lr28* which eliminates the possibility of association of *Lr28* with *scaffold163140-5_613860*.Further, *scaffold163140-5_613860* was also mapped in the vicinity of a recently reported MTA (*AX-95106749*) for adult plant LR resistance (Vikas et al., 2022). Since, *scaffold163140-5_613860* was identified for seedling resistance to LR, it is unlikely that this MTA represents *AX-95106749*. Similarly, *scaffold56230_857271, scaffold151621_1347043*, and *scaffold33098_3932746*, identified on chromosomes 2B, 3B, and 5B represent a novel locus associated with TNBJS and MNPSD respectively, as no previously reported *Lr* gene has been detected in this region.

Next, we selected 17 significant MTAs based on LD decay rate of less than 1Mbp to identify candidate genes with putative role in disease resistance (Supplementary Tables S8 and S9). The putative candidate genes belonged to different disease resistance encoding gene families such as leucine-rich repeats receptor-like kinases (LRR-RLKs), nucleotide-binding site leucine-rich repeats (NBS-LRRs), serine/threonine-protein phosphatase domain containing proteins, ABC transporters, zinc finger proteins, and others (Supplementary Table S9). Zinc finger domains have been identified in various disease resistance genes from several crops, indicating their significant role in conferring host-plant resistance (Epple et al., 2003; Ciftci-Yilmaz and Mittler, 2008; Emerson and Thomas, 2009). Further, serine/threonine-protein phosphatase domain containing proteins are known to be involved in regulation of plant defense and stress responses (País et al., 2009; Máthé et al., 2019). LRR-RLKs and ABC transporters have known to be involved in a wide variety of developmental and defense-related processes (Torii, 2004; Krattinger et al., 2009; Kang et al., 2011). Additionally, a few more genes encoding putative proteins of interest including plant defensins, germin-like proteins, E3 ubiquitin-protein ligase, cytochrome P450 family protein, F-box proteins (FBPs), and others were identified (Supplementary Tables S8 and S9). These identified genes could serve as valuable information in future gene cloning efforts of these genomic regions.

In conclusion, we identified valuable sources of LR resistance against multiple *Pt* races. The SNP markers reported as associated with resistance can facilitate the deployment of these QTLs through marker-assisted selection in breeding programmes.

## Acknowledgments

The authors would like to thank Julie Hochhalter for providing the greenhouse resources at Dalrymple Research Greenhouse complex at Fargo, ND.

## Author Contributions

UG and SKS conceptualized the experiment and designed the methodology; SK, and UG performed the investigation; SK and HG performed the data curation, data analysis, and visualization; SK and HG performed the software implementation; SK, HG, SKS, and UG wrote the original manuscript; SK, HG, JK, RG, SKS, and UG contributed to the interpretation of results; All the authors contributed to manuscript revision and approved the final manuscript.

## Ethical approval

The authors declare that the experiments comply with the current laws of the country.

## Supplementary Information (SI)

The Supplementary information cited in the manuscript can be found in an attached PDF document.

## Figures

Supplementary Figure S1. Correlation analysis among the phenotypic data of 365 wheat accessions evaluated for their reaction to *P. triticina* races TDBJQ, TBBGS, MNPSD, and TNBJS.

Supplementary Figure S2. Scatterplot based on the first two components (PC1 and PC2) from PCA for (**A**) growth habit and (**B**) geographic origin.

Supplementary Figure S3. Intra-chromosomal linkage disequilibrium in the diversity panel for (**A**) whole genome and, individually in the (**B**) A, (**C**) B, and (**D**) D sub-genomes.

Supplementary Figure S4. Box-cox transformations performed for *Pt* races TDBJQ, TBBGS, MNPSD, and TNBJS to normalize the data.

