## Supplementary figures and images for "Identification of leaf rust resistance loci in a geographically diverse panel of wheat using genome-wide association analysis"

### Supplementary Figure S1

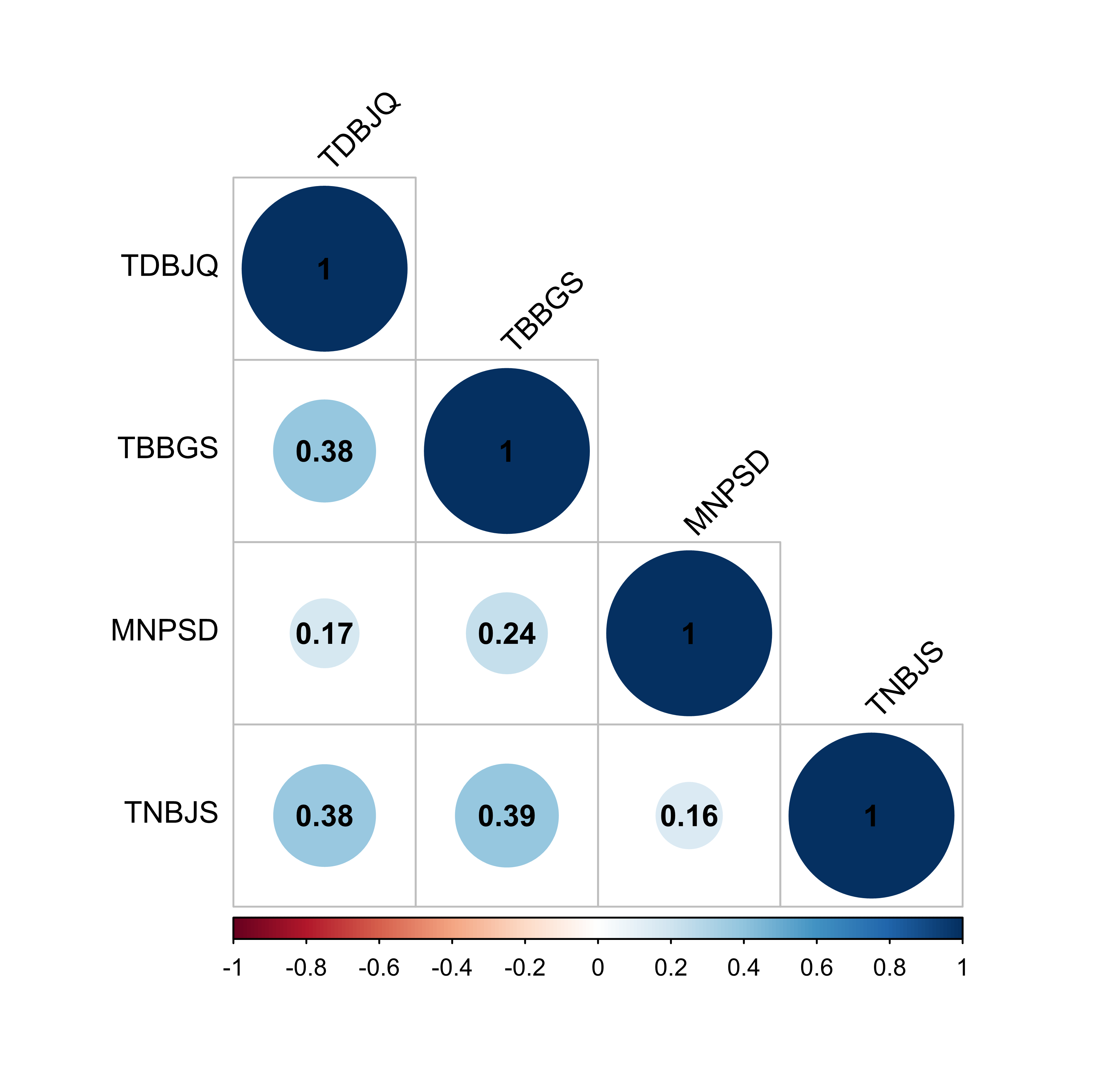

### Supplementary Figure S2

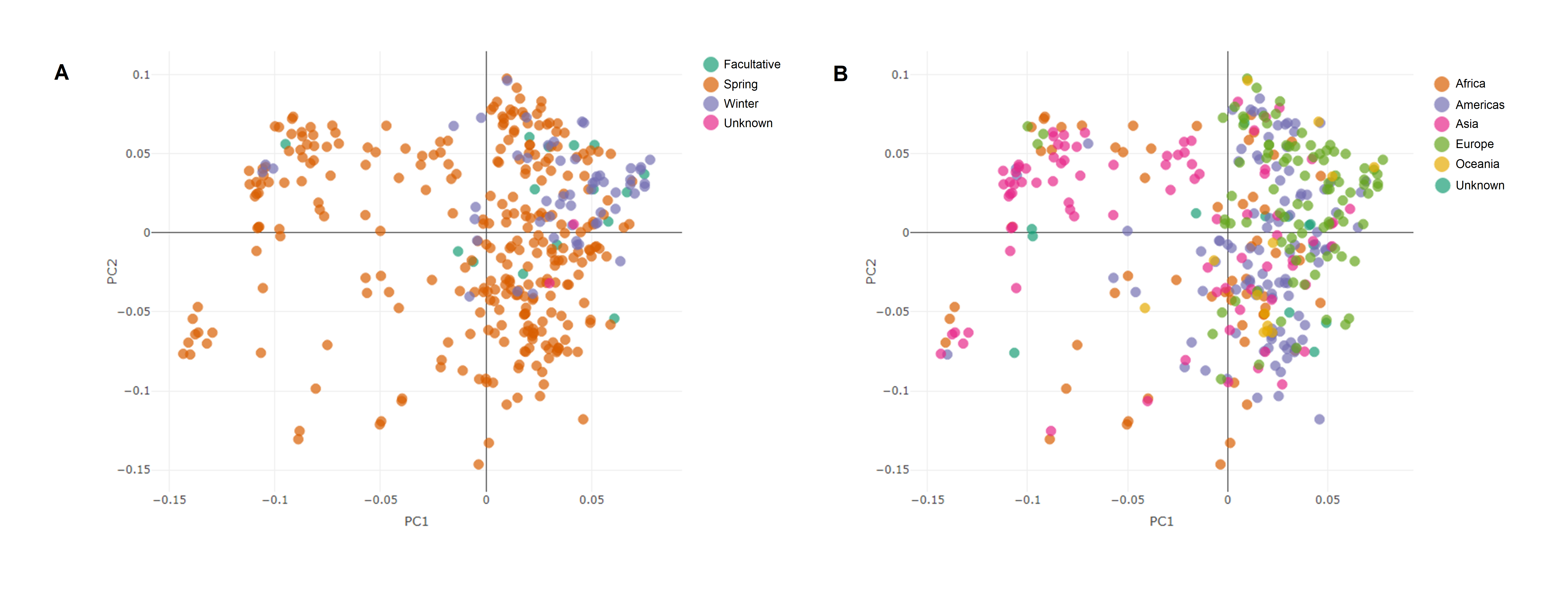

### Supplementary Figure S3

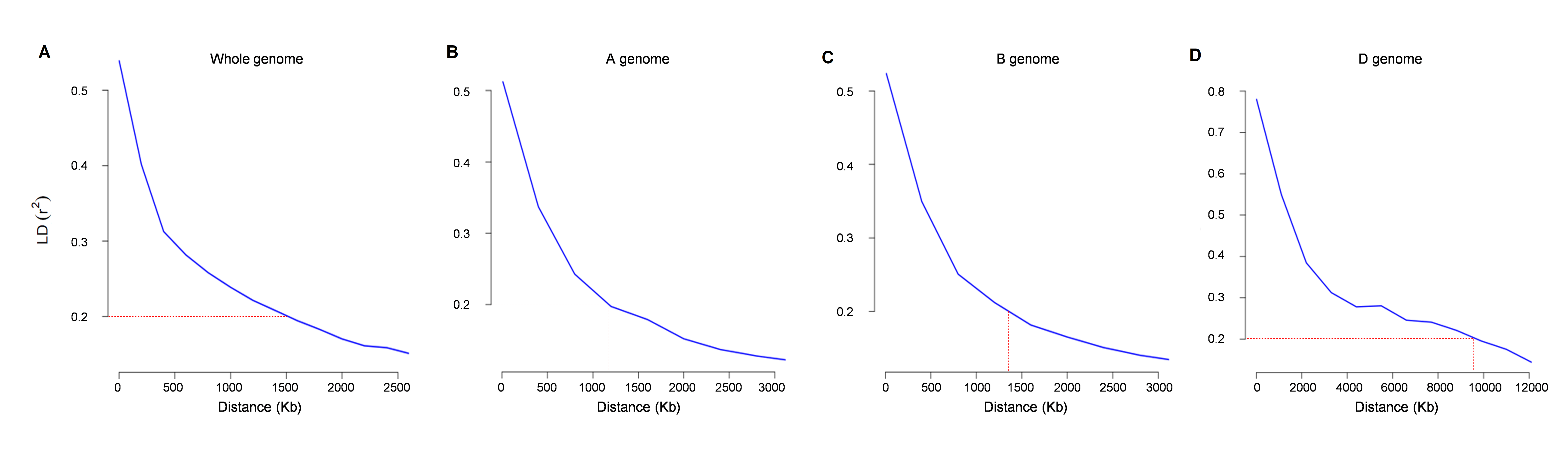

### Supplementary Figure S4

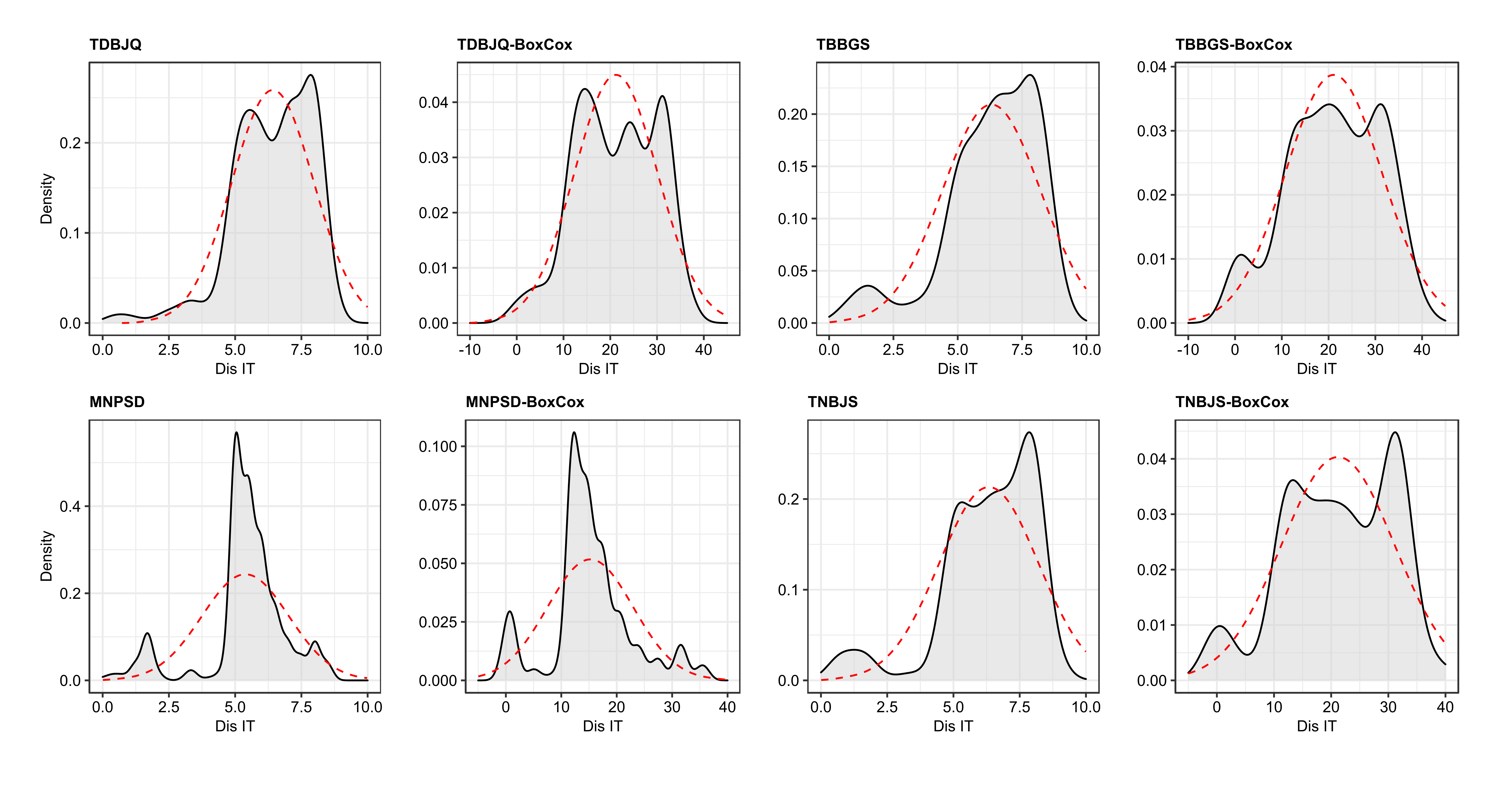
